# Nicotinamide-*N*-methyltransferase is essential for SAM and 1-methylnicotinamide homeostasis in the AML12 hepatocyte cell line

**DOI:** 10.1101/2022.09.25.509348

**Authors:** Mayuko Yoda, Rin Mizuno, Yoshihiro Izumi, Masatomo Takahashi, Takeshi Bamba, Shinpei Kawaoka

## Abstract

Nicotinamide-*N*-methyltransferase (NNMT) is an enzyme that consumes *S*-adenosyl-methionine (SAM) and nicotinamide (NAM) to produce *S*-adenosyl-homocysteine (SAH) and 1-methylnicotinamide (MNAM). How much NNMT contributes to the quantity regulation of these four metabolites depends on whether NNMT is a major consumer or producer of these metabolites, which varies among various cellular contexts. Yet, whether NNMT critically regulates these metabolites in the AML12 hepatocyte cell line has been unexplored. To address this, we knock down *Nnmt* in AML12 cells and investigate the effects of *Nnmt* RNAi on metabolism and gene expression. We find that *Nnmt* RNAi accumulates SAM and SAH, whereas it reduces MNAM with NAM being unaltered. These results indicate that NNMT is a significant consumer of SAM and critical for MNAM production in this cell line. Moreover, transcriptome analyses reveal that altered SAM and MNAM homeostasis is accompanied by various detrimental molecular phenotypes, as exemplified by the down-regulations of lipogenic genes such as *Srebf1*. Consistent with this, oil-red O-staining experiments demonstrate the decrease of total lipids upon *Nnmt* RNAi. These results suggest that NNMT maintains proper SAM and MNAM homeostasis, providing an additional example where NNMT plays a critical role in regulating SAM and MNAM metabolism.

## Introduction

Nicotinamide-*N*-methyltransferase (NNMT) transfers a methyl group from *S*-adenosyl-methionine (SAM) to nicotinamide (NAM) and produces *S*-adenosyl-homocysteine (SAH) and 1-methylnicotinamide (MNAM) (*1*). SAM is a methyl donor that contributes to various methylation events in the cytoplasm and nucleus (*2–4*). In addition, recent studies demonstrate that MNAM retains biological activities such as anti-inflammation (*5–10*). Via regulating the NNMT-related metabolites, NNMT plays a critical role in a series of phenomena such as energy metabolism (*1, 5, 7, 8, 11–18*).

How much NMNT contributes to the quantity of SAM, NAM, MNAM, and SAH is context-dependent: it depends on how many other enzymes are involved in the metabolism of these four metabolites in a certain cellular context. For example, suppression of NNMT results in the loss of MNAM in all cell types reported so far (*1, 5, 7, 10–13*). This simplicity is attributed to the fact that only NNMT produces MNAM in worms, mice, and humans. On the other hand, the homeostasis of SAM and NAM is more complex, as many other enzymes consume and produce these two metabolites. In the case of murine livers, deletion of NNMT does not accumulate SAM and NAM, suggesting that NNMT is not a significant consumer for them in the liver. Intriguingly, when GNMT, the major consumer for SAM in the liver, is reduced, suppression of NNMT leads to the accumulation of SAM (i.e., the contribution of NNMT to SAM homeostasis is increased upon GNMT suppression) (*7, 13*). These studies exemplify that NNMT context-dependently contributes to SAM homeostasis. As such, the effects of altered NNMT metabolism on cellular homeostasis should differ among contexts. In this regard, investigation of the roles of NNMT in various cellular contexts is essential to deepen our understanding of the significance of NNMT-dependent metabolism in living cells.

In the current study, we examine the roles of NNMT in metabolism and gene expression in the AML12 hepatocyte cell line. AML12 is one of the commonly used murine hepatocyte cell lines established from mice overexpressing human transforming growth factor-alpha (TGF-a) (*19*). We show that NNMT regulates SAM and MNAM homeostasis, and maintains proper gene expression and lipid metabolism in this hepatocyte cell line. This study provides one more example where NNMT significantly contributes to SAM and MNAM homeostasis.

## Results

### RNAi efficiently reduces *Nnmt* expression in AML12 cells

To address the roles of NNMT in the AML12 hepatocyte cell line, we designed small interfering RNAs (siRNAs) against this gene (*siNnmt*) (Fig. 1a). siRNAs targeting luciferase (*siLuc*) were used as a control. We treated AML12 cells with either si*Luc* or *siNnmt* for 48 hours and measured the expression of *Nnmt* using quantitative reverse transcription PCR (qRT-PCR). Our data demonstrated that si*Nnmt* effectively reduced the *Nnmt* mRNA levels by 91% (Fig. 1b). These results indicated that the si*Nnmt* experiments enable us to evaluate the roles of NNMT in metabolism and gene expression in this hepatocyte cell line.

**Figure 1:**
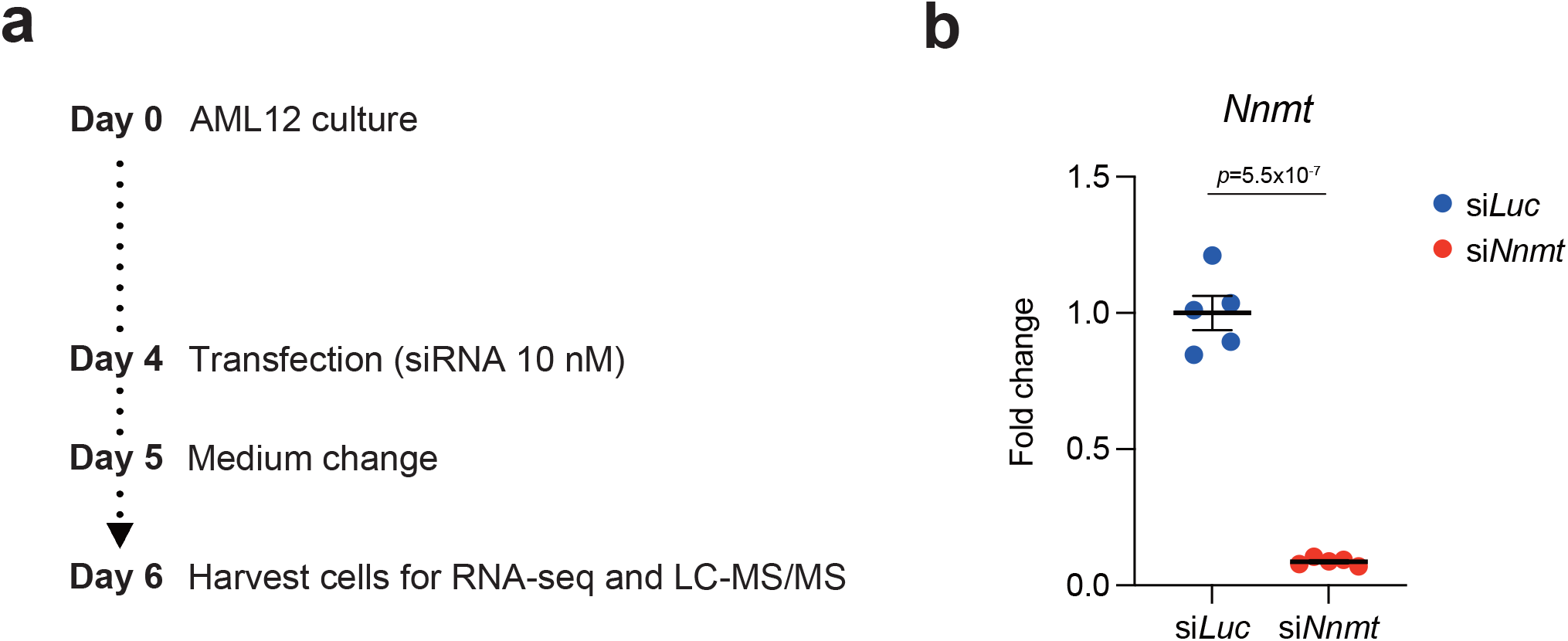
*Nnmt* RNAi in the AML12 hepatocyte cell line. **(a)** The scheme of the *Nnmt* RNAi experiments in AML12 cells **(b)** qPCR analysis for *Nnmt* in AML12 cells treated with *siLuc* or si*Nnmt*. Averaged fold change data normalized to the si*Luc* group are presented as the mean ±SEM. *n* = 5. The *p* values are shown (unpaired two-tailed Student’s *t*-test).

### *Nnmt* RNAi accumulates SAM and decreases MNAM in AML12 cells

We then analyzed the amount of SAM, NAM, SAH, and MNAM in AML12 cells treated with si*Luc* or si*Nnmt* with the aid of liquid chromatography coupled with tandem mass spectrometry (LC-MS/MS). We found that *Nnmt* RNAi reduces MNAM (Fig. 2b and Table S1). This was consistent with many other studies that demonstrate the essential role of NNMT in MNAM production (*1, 5, 7, 10–13*). The significant decrease in MNAM validated the efficiency of *Nnmt* RNAi in AML12 cells. On the other hand, *Nnmt* RNAi did not affect NAM, suggesting that NNMT does not have a significant impact on maintaining the quantity homeostasis of this metabolite in AML12 cells (Fig. 2c).

**Figure 2:**
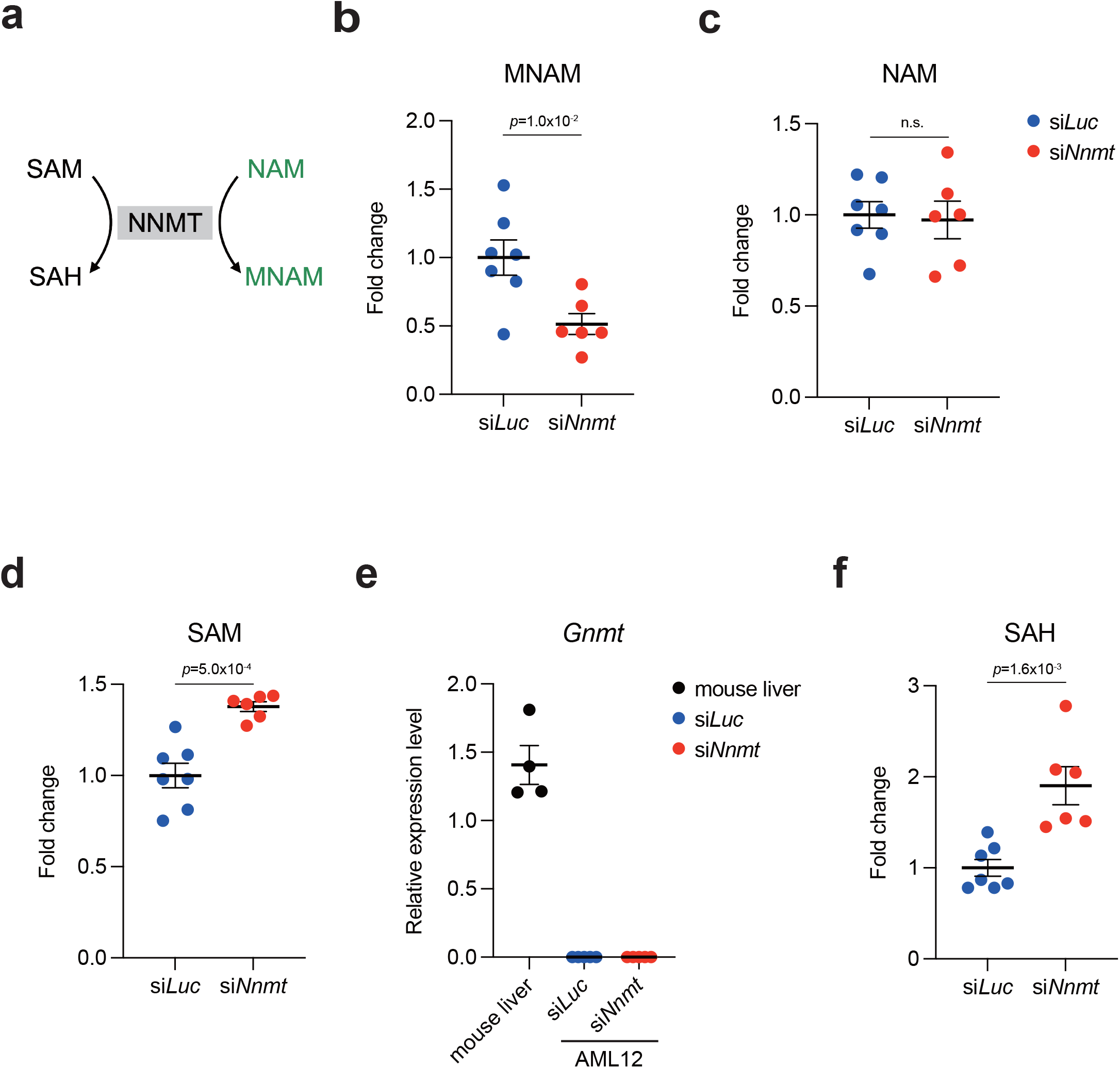
*Nnmt* RNAi accumulates SAM and decreases MNAM in AML12 cells. **(a)** The biochemical reaction catalyzed by NNMT. **(b)** LC-MS/MS analysis for MNAM. **(c)** LC-MS/MS analysis for NAM. **(d)** LC-MS/MS analysis for SAM. **(e)** qPCR analysis for *Gnmt* in the livers and AML12 cells. Relative expression normalized to *B2m* are shown as the mean ±SEM. *n* = 4 for the livers and *n* = 5 for AML12 cells. **(f)** LC-MS/MS analysis for SAH. In **(b)**-**(d)** and **(f)**, averaged fold change data normalized to the si*Luc* group are presented as the mean ± SEM. The *p* values are shown (unpaired two-tailed Student’s *t*-test. *n* = 7 for the si*Luc* group and *n* = 6 for the si*Nnmt* group).

Notably, we found that *siNnmt* elevated SAM in AML12 cells (Fig. 2d). This was an intriguing contrast to the published data in which suppression of *Nnmt* does not increase SAM in the murine liver (*7, 13*). As described earlier, the most prominent consumer of SAM in the liver is glycine-*N*-methyltransferase (GNMT), whose knockout elevates SAM (*13, 20*). Hong et al. also have shown that *Nnmt* knockdown in mouse primary hepatocytes accumulate SAM when *Gnmt* is knocked down simultaneously (*13*). To this end, we compared *Gnmt* expression in the liver and AML12 cells, finding that *Gnmt* is relatively less abundant in AML12 cells when compared to the liver (Fig. 2e). Curiously, SAH was also elevated upon *Nnmt* RNAi (Fig. 2f). This implied that part of SAM accumulated by *Nnmt* RNAi was then metabolized to SAH by other methyltransferases. Collectively, in this cell line, NNMT significantly contributed to SAM homeostasis, presumably due to the low expression of *Gnmt* (see also Discussion regarding possible involvement of another metabolic pathway). These data provide additional evidence that the contribution of NNMT in SAM homeostasis varies depending on cellular contexts.

### *Nnmt* RNAi disrupts global gene expression and lipogenesis in AML12 cells

Given the critical importance of NNMT in SAM and MNAM metabolism in AML12 cells, we wanted to investigate if the altered SAM and MNAM homeostasis affects global gene expression in this cell line. To this end, we performed transcriptome analyses. As shown in Fig. 3a, *Nnmt* RNAi resulted in down-regulation of 747 genes and up-regulation of 65 genes (> 2-fold change and *q* value < 0.05; Table S2), suggesting that the NNMT-dependent metabolism is important in maintaining proper gene expression in AML12 cells. Such crucial roles of NNMT were exemplified by the reduction of *Albumin (Alb*), the marker gene for AML12 hepatocytes (Fig. 3b). Gene ontology analyses demonstrated that the down-regulated genes represent dampened “lipid metabolic process” in this cell line (Fig. 3c). Genes including *diacylglycerol acyltransferase 2 (Dgat2)* (*21*) and *sterol regulatory element binding transcription factor 1 (Srebf1)* (*22*) were severely reduced upon *Nnmt* RNAi (Fig. 3d). It has been shown that these genes are involved in lipogenesis, implicating a role of NNMT in lipogenesis in AML12 cells. To further test this hypothesis, we performed oil-red O-staining against AML12 cells treated with either *siLuc* or si*Nnmt*. We quantified total lipid levels using a spectrophotometer (OD = 540 nm), finding that total lipids were reduced upon *Nnmt* RNAi (Fig. 3e). Although it is currently unknown how NNMT facilitates lipogenesis, our study demonstrated a significant contribution of this enzyme in the lipogenesis pathway in this cell line.

**Figure 3:**
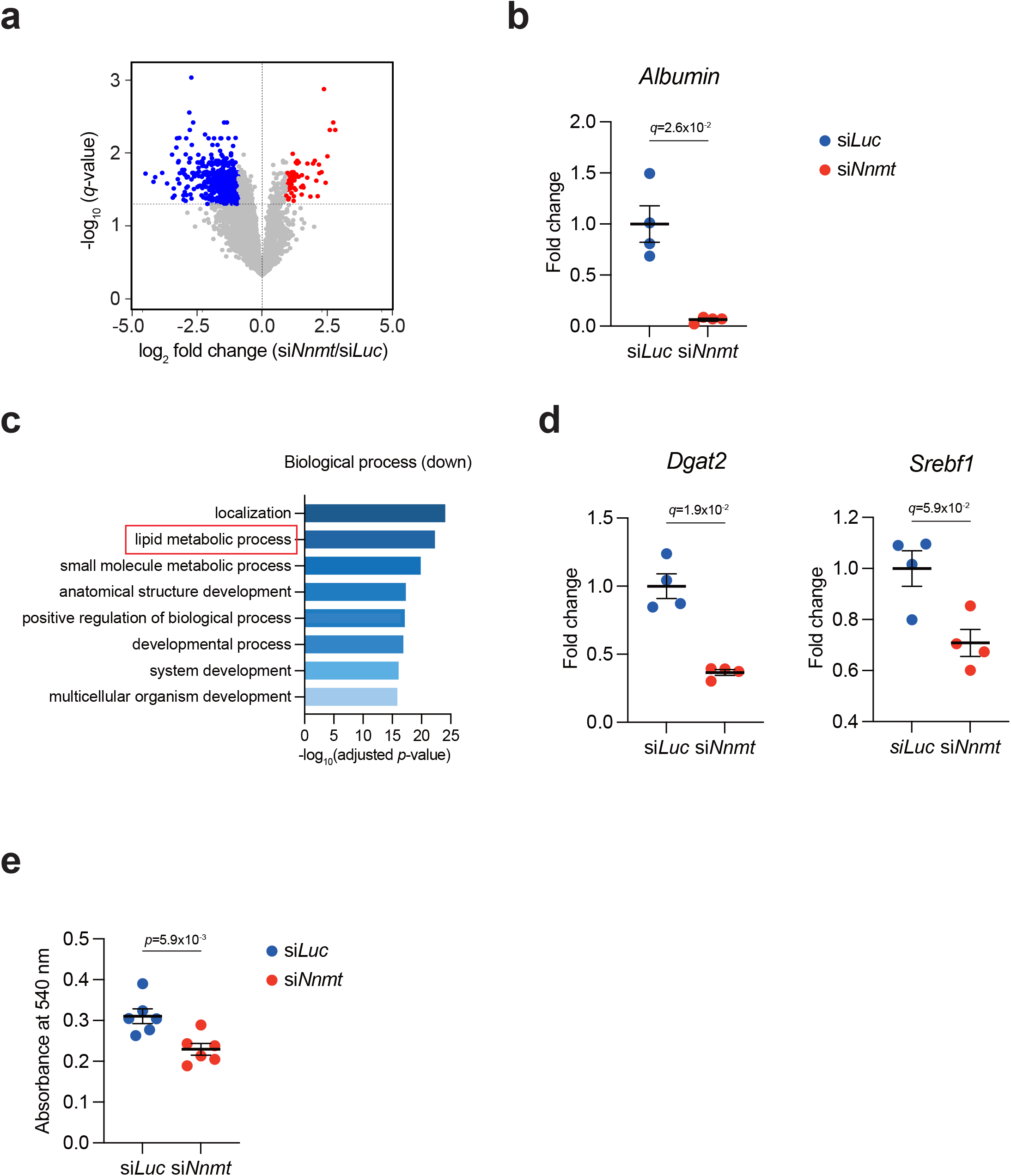
*Nnmt* RNAi impairs global gene expression and lipogenesis in AML12 cells. **(a)** RNA-seq experiments for AML12 cells treated with si*Luc* or si*Nnmt*. A volcano plot (log_2_-fold average (*siNnmt/siLuc*) versus −log_10_ (*q* value)) is shown. Genes showing more than 2-fold change with *q* < 0.05 are highlighted. *n* = 4. **(b)** RNA-seq results of representative down-regulated gene *Albumin*. **(c)** Gene ontology analysis (g:Profiler) for genes that are down-regulated by *Nnmt* RNAi. Adjusted enrichment *p* values obtained from g:Profiler are shown. **(d)** RNA-seq results of representative down-regulated genes involved in lipogenesis (*Dgat2* and *Srebf1*). **(e)** Oil-red O staining. Total lipids are measured as the dye extraction signals. *n* = 6. In **(b)** and **(d)**, averaged fold change data normalized to the si*Luc* group are presented as the mean ± SEM. The *q* values are shown.

## Discussion

In the current study, we examined the roles of NNMT in metabolism and gene expression in the AML12 hepatocytes cell line. We found that NNMT is critical to maintaining SAM and MNAM homeostasis and contributes to lipogenesis in this cell line.

As discussed earlier, the degree of contribution of NNMT to SAM homeostasis differs depending on cell types. We previously demonstrated that, in the murine livers, deletion of *Nnmt* does not alter the steady-state amount of SAM in healthy conditions (*7*) (Fig. 4). We reasoned that this was because GNMT is the major consumer of SAM in the liver in vivo and because the trigonelline pathway might have received excess methyl groups from SAM (Fig. 4). Trigonelline is a methylated form of nicotinic acids (*23*). Interestingly, the murine livers do not have trigonelline-producing activity, suggesting that trigonelline is synthesized with the help of, for example, microbiomes (*7, 23*). Consistent with this assumption, we found that trigonelline was undetectable in AML12 cells that must have been free from microbiomes (Table S1). Hence, in contrast to living mice, AML12 cells seemingly lack the trigonelline pathway that could receive methyl groups from SAM. In addition, the expression of *Gnmt* is low in this cell line (Fig. 2e). We suggest that these biological contexts establish NNMT as a crucial regulator of SAM homeostasis in AML12 cells. In summary, this study provides additional evidence that the methyl-donor balance is maintained in context-dependent manners.

**Figure 4:**
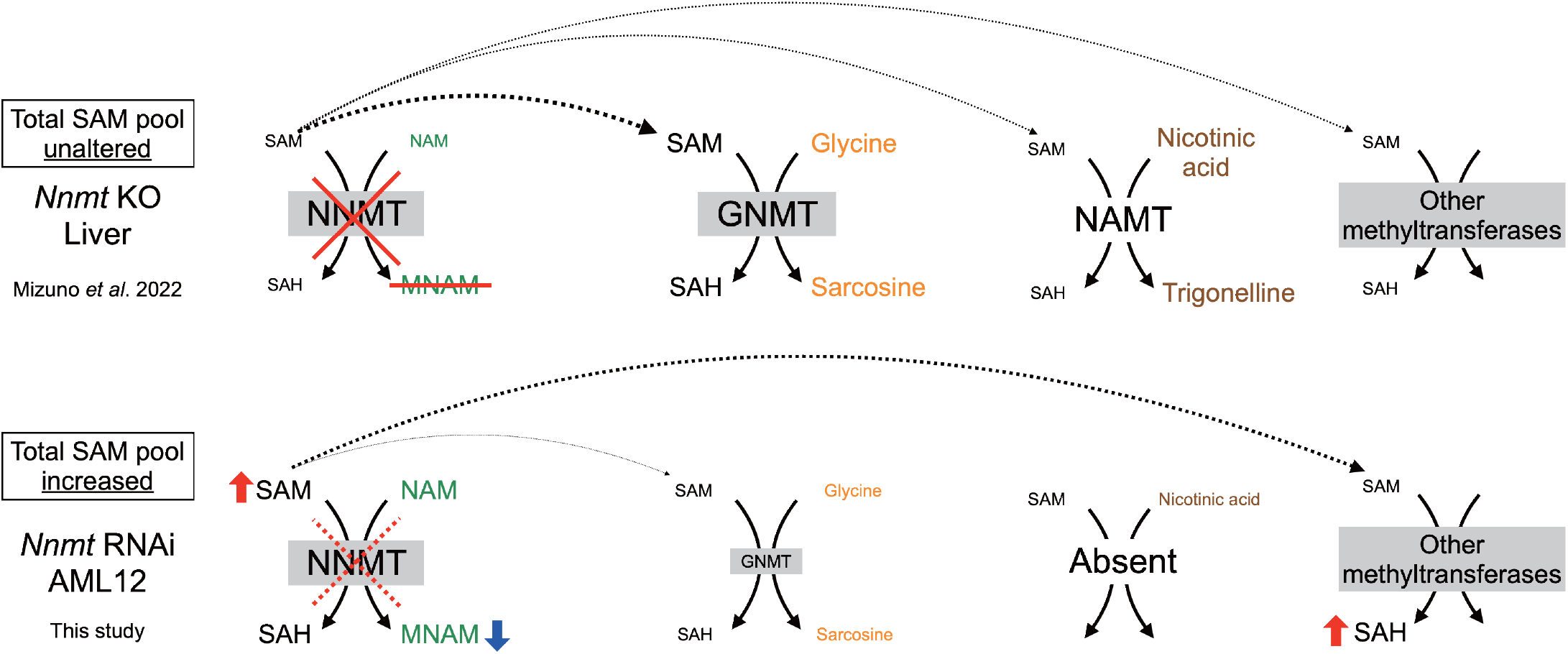
The summary of this study. The contribution of NNMT in the maintenance of SAM is context-dependent. In the liver, SAM homeostasis is maintained even in the absence of NNMT. This is likely owing to the strong contribution of GNMT in SAM homeostasis. In addition, nicotinic acid methyltransferase (NAMT), which is derived most likely from the microbiome (*7, 23*), seemingly contributes to SAM homeostasis in the liver. Compared to the liver, the expression of *Gnmt* in AML12 cells is relatively low. In addition, AML12 cells lack the NAMT-trigonelline pathway. In these conditions, NNMT plays a critical role in SAM homeostasis.

In line with the critical roles in SAM and MNAM homeostasis (Fig. 2), we found that NNMT significantly contributes to global gene expression in AML12 cells (Fig. 3). Most importantly, *Nnmt* RNAi impaired lipogenesis in this cell line (Fig. 3e). This finding is consistent with the recent study reported by Song and colleagues: they show that *Nnmt* knockdown and the administration of NNMT inhibitors suppress lipogenesis in a mouse model of fatty liver diseases and AML12 cells (*24*). Thus, our study additionally solidifies that NNMT positively regulates lipogenesis in hepatocytes. Whether and how SAM and MNAM metabolism are involved in lipogenesis awaits further examination.

NNMT is a cytoplasmic enzyme. It is thus unlikely that NNMT protein control gene expression directly. Yet, its substrate SAM and product MNAM could contribute to gene expression. We expect that the excess SAM and the reduction in MNAM at least partly should account for gene expression changes caused by *Nnmt* RNAi. However, it is unlikely that all gene expression changes detected in *Nnmt*-knock-down AML12 cells are directly owing to such metabolic changes. Based on this discussion, we assume that our datasets include alterations secondarily caused by SAM- and MNAM-dependent changes for their target genes (i.e., secondary effects of *Nnmt* RNAi). Further studies are required to discriminate primary and secondary gene expression changes by *Nnmt* deficiency and to reveal how SAM and MNAM regulate gene expression.

Collectively, our data showed that NNMT is critical for maintaining global gene expression and lipogenesis in this hepatocyte cell line. These data deepened our understanding of how NNMT regulates metabolism and gene expression in various cellular contexts.

## Methods

### Cell culture

AML12 cells were obtained from American Type Culture Collection (ATCC, VA, USA: CRL-2254^™^) and cultured in DMEM/Ham’s F-12 (nacalai tesque, Kyoto, Japan) containing 10% fetal bovine serum, 40 ng/ml dexamethasone (nacalai tesque), and 1×Insulin-Transferrin-Selenium (Gibco, CA, USA) in a 5% CO2 tissue culture incubator at 37°C.

### siRNA transfection

AML12 cells were seeded at 1×10^5^ cells/well in a 6-well plate and transfected with siRNA mixture (f.c. 10 nM) using Lipofectamine 2000 (Invitrogen, MA, USA) according to the manufacturer’s instructions. At 24 hours after transfection, the medium was exchanged with new DMEM/Ham’s F-12 containing 10% fetal bovine serum, 40 ng/ml Dexamethasone (nacalai tesque), and 1×Insulin-Transferrin-Selenium (Gibco). The treated cells were further cultured for 24 hours and then harvested. The sequences of siRNAs are as follows:

si*Luc*_S (CUUACGCUGAGUACUUCGAUU)

si*Luc*_AS (UCGAAGUACTCAGCGUAAGUU)

si*Nnmt*_S (AGGCCUGCUGGUUCAUUUCUU)

si*Nnmt*_AS (GAAAUGAACCAGCAGGCCUUU)

### Quantitative reverse transcription PCR

Total RNAs were extracted from AML12 cells and mouse livers using RNeasy Mini Kit (Qiagen, Venlo, Nederland) according to the manufacturer’s protocol. cDNA was synthesized with Transcriptor First Strand cDNA synthesis kit (Roche, Basel, Switzerland). qPCR experiments were performed using QuantStudio 5 Real-time PCR system (Thermo Fisher Scientific, MA, USA) and SYBR Green Master Mix (Roche). *B2m* was used as an internal control. The primers used in these experiments are listed as follows:

*Nnmt*_F (GAACCAGGAGCCTTTGACTG), *Nnmt*_R (GATTGCACGCCTCAACTTCT),

*Gnmt*_F (AGCCCACATGGTAACCCTGG), *Gnmt*_R (TGAAGTCACCCAGGACGCTG),

*B2m*_F (GCTCGGTGACCCTGGTCTTT), *B2m*_R (AATGTGAGGCGGGTGGAACT).

### Liquid chromatography coupled with tandem mass spectrometry

Metabolites from AML12 cells were extracted using the Blight and Dyer’s method (*25*) with some modifications. Briefly, each sample was mixed with 1 ml of cold methanol containing 10-camphorsulfonic acid (1.0 nmol or 1.5 nmol) as internal standard (IS) for mass spectrometrybased metabolomic analysis. The samples were vigorously mixed by vortexing for 1 min followed by 5 min of sonication. The extracts were then centrifuged at 16,000 ×*g* for 5 min at 4°C, and the resultant supernatant was collected. After mixing supernatant with chloroform and water (methanol:chloroform:water = 5:5:4), the aqueous and organic layers were separated by vortexing and subsequent centrifugation at 16,000 × *g* and 4°C for 5 min. The aqueous (upper) layer was transferred into a clean tube. After the aqueous layer extracts were evaporated under vacuum, the dried extracts were stored at −80°C until the analysis of hydrophilic metabolites. Prior to analysis, the dried aqueous layer was reconstituted in 30 to 50 μl of water. Liquid chromatography tandem mass spectrometry (LC/MS/MS) methods for hydrophilic metabolite analysis were employed as described previously (*26, 27*). Cationic polar metabolites were analyzed via liquid chromatography (Nexera X3 UHPLC system, Shimadzu, Kyoto, Japan) with a Discovery HS F5 column (2.1 mm i.d. × 150 mm, 3 μm particle size, Merck) coupled with a LCMS-8060NX, triple quadrupole mass spectrometer (Shimadzu) and via liquid chromatography (Nexera X2 UHPLC system, Shimadzu) with a Discovery HS F5 column (2.1 mm i.d. × 150 mm, 3 μm particle size, Merck) coupled with a Q Exactive instrument. The analytical platform for hydrophilic metabolite analysis was controlled using LabSolutions (version 5.80) and LabSolutions Insight (version 3.80) (Shimadzu). The quantitative content of the hydrophilic metabolites was calculated using peak area relative to the IS.

### Transcriptome analysis

Total RNAs were extracted from AML12 cells as described above with RNase-Free DNase Set (Qiagen). RNA-seq libraries were generated using the NEBNext Globin&rRNA depletion kit and the NEBNext UltraII Directional RNA Library prep kit according to the manufacturer’s instructions (New England Biolabs, MA, USA). Sequencing experiments were performed with NextSeq 500 (Illumina; High Output Kit v2.5, 75 Cycles). The obtained reads were mapped to the mouse genome grcm38 and processed using fastp (removing reads with < Q30), Hisat2, Samtools, and featureCounts (*28–31*). The obtained expression matrix with TPM scores is shown in Supplementary Data 2. The volcano plot was depicted using ggplot2 to visualize differentially expresses genes (https://ggplot2.tidyverse.org/index.html). Differentially expressed genes were further subjected to gene ontology analyses using g:Profiler (*32*).

### Oil-red O staining

To measure total lipids, we stained AML12 cells using the Lipid Assay Kit (COSMO BIO, Tokyo, Japan) according to the manufacturer’s instructions. Briefly, AML12 cells were treated with either *siLuc* or si*Nnmt* for 48 hours. The cells were then washed with PBS and fixed with 10% formalin at room temperature overnight. The fixed AML12 cells were washed three times with distilled water and stained with oil-red O at room temperature for 15 min. Following staining, the cells were washed three times with distilled water and dried at room temperature overnight. The dye extractions from these cells were quantified by measuring 540 nm by a spectrophotometer Multiskan GO (Thermo Fisher Scientific).

### Statistics and Data visualization

GraphPad Prism Software was used to analyze data. Data were displayed as mean ± SEM. Student’s *t* test was performed to analyze the statistical significance between groups, and *p* value < 0.05 was considered statistically significant. In RNA-seq analyses, *q* value was calculated using the Storey’s method (https://www.bioconductor.org/packages/release/bioc/html/qvalue.html).

## Data availability

RNA-seq data obtained in this study are available from DNA Databank of Japan under the accession number of DRA014854.

## Acknowledgements

This work was supported by JSPS KAKENHI (16H06279, 18K15409, 18H04810, 20H03451, 20H04842, and 22H04925; S.K), JST FOREST (20351876; S.K), JST Moonshot (JPMJMS2011-61; S.K), Caravel, Co., Ltd (S.K), Ono Medical Research Foundation (S.K), Takeda Science Foundation, The Uehara memorial foundation (S.K), Chubei Ito foundation (S.K) and Japan Foundation for Applied Enzymology (S.K). We thank Dr. Takefumi Kondo and Yukari Sando for their support in the transcriptome analyses.

## Author contributions

M.Y. performed experiments, analyzed data, constructed figures, and wrote the paper. R.M. performed experiments and analyzed data. Y.I., M.T., M.N., and T.B. performed metabolites measurements. S.K. conceived and supervised this study, analyzed data, and wrote the manuscript.

## Competing interests

The authors declare no competing interests exist in this study.

## Notes

### Competing Interest Statement

The authors have declared no competing interest.

### Summary of Updates

Changed "Distribution/Reuse Options"

## References

(1) Pissios, P. (2017) Nicotinamide N-Methyltransferase: More Than a Vitamin B3 Clearance Enzyme. Trends Endocrinol Metab. 28, 340–353

(2) Lu, S.C. (2000) S-Adenosylmethionine. Int J Biochem Cell Biol. 32, 391–395

(3) Ulrey, C.L., Liu, L., Andrews, L.G., and Tollefsbol, T.O. (2005) The impact of metabolism on DNA methylation. Hum Mol Genet. 14 Spec No 1, R139–147

(4) Luo, M. (2012) Current chemical biology approaches to interrogate protein methyltransferases. ACS Chem Biol. 7, 443–463

(5) Hong, S., Moreno-Navarrete, J.M., Wei, X., Kikukawa, Y., Tzameli, I., Prasad, D., Lee, Y., Asara, J.M., Fernandez-Real, J.M., Maratos-Flier, E., and Pissios, P. (2015) Nicotinamide N-methyltransferase regulates hepatic nutrient metabolism through Sirt1 protein stabilization. Nat Med. 21, 887–894

(6) Nejabati, H.R., Mihanfar, A., Pezeshkian, M., Fattahi, A., Latifi, Z., Safaie, N., Valiloo, M., Jodati, A.R., and Nouri, M. (2018) N1-methylnicotinamide (MNAM) as a guardian of cardiovascular system. J Cell Physiol. 233, 6386–6394

(7) Mizuno, R., Hojo, H., Takahashi, M., Kashio, S., Enya, S., Nakao, M., Konishi, R., Yoda, M., Harata, A., Hamanishi, J., Kawamoto, H., Mandai, M., Suzuki, Y., Miura, M., Bamba, T., Izumi, Y., and Kawaoka, S. (2022) Remote solid cancers rewire hepatic nitrogen metabolism via host nicotinamide-N-methyltransferase. Nat Commun. 13, 3346

(8) Kraus, D., Yang, Q., Kong, D., Banks, A.S., Zhang, L., Rodgers, J.T., Pirinen, E., Pulinilkunnil, T.C., Gong, F., Wang, Y.C., Cen, Y., Sauve, A.A., Asara, J.M., Peroni, O. D., Monia, B.P., Bhanot, S., Alhonen, L., Puigserver, P., and Kahn, B.B. (2014) Nicotinamide N-methyltransferase knockdown protects against diet-induced obesity. Nature. 508, 258–262

(9) Kilgour, M.K., MacPherson, S., Zacharias, L.G., Ellis, A.E., Sheldon, R.D., Liu, E.Y., Keyes, S., Pauly, B., Carleton, G., Allard, B., Smazynski, J., Williams, K.S., Watson, P. H., Stagg, J., Nelson, B.H., DeBerardinis, R.J., Jones, R.G., Hamilton, P.T., and Lum, J.J. (2021) 1-Methylnicotinamide is an immune regulatory metabolite in human ovarian cancer. Sci Adv. 7,

(10) Schmeisser, K., Mansfeld, J., Kuhlow, D., Weimer, S., Priebe, S., Heiland, I., Birringer, M., Groth, M., Segref, A., Kanfi, Y., Price, N.L., Schmeisser, S., Schuster, S., Pfeiffer, A.F., Guthke, R., Platzer, M., Hoppe, T., Cohen, H.Y., Zarse, K., Sinclair, D.A., and Ristow, M. (2013) Role of sirtuins in lifespan regulation is linked to methylation of nicotinamide. Nat Chem Biol. 9, 693–700

(11) Ulanovskaya, O.A., Zuhl, A.M., and Cravatt, B.F. (2013) NNMT promotes epigenetic remodeling in cancer by creating a metabolic methylation sink. Nat Chem Biol. 9, 300–306

(12) Sperber, H., Mathieu, J., Wang, Y., Ferreccio, A., Hesson, J., Xu, Z., Fischer, K.A., Devi, A., Detraux, D., Gu, H., Battle, S.L., Showalter, M., Valensisi, C., Bielas, J.H., Ericson, N.G., Margaretha, L., Robitaille, A.M., Margineantu, D., Fiehn, O., Hockenbery, D., Blau, C.A., Raftery, D., Margolin, A.A., Hawkins, R.D., Moon, R.T., Ware, C.B., and Ruohola-Baker, H. (2015) The metabolome regulates the epigenetic landscape during naive-to-primed human embryonic stem cell transition. Nat Cell Biol. 17, 1523–1535

(13) Hong, S., Zhai, B., and Pissios, P. (2018) Nicotinamide N-Methyltransferase Interacts with Enzymes of the Methionine Cycle and Regulates Methyl Donor Metabolism. Biochemistry. 57, 5775–5779

(14) Wang, Y., Zeng, J., Wu, W., Xie, S., Yu, H., Li, G., Zhu, T., Li, F., Lu, J., Wang, G.Y., Xie, X., and Zhang, J. (2019) Nicotinamide N-methyltransferase enhances chemoresistance in breast cancer through SIRT1 protein stabilization. Breast Cancer Res. 21, 64

(15) Li, D., Yi, C., Huang, H., Li, J., and Hong, S. (2022) Hepatocyte specific depletion of Nnmt protects the mice from non-alcoholic steatohepatitis. J Hepatol.

(16) Wang, W., Yang, C., Wang, T., and Deng, H. (2022) Complex roles of nicotinamide N-methyltransferase in cancer progression. Cell Death Dis. 13, 267

(17) Ogawa, M., Tanaka, A., Namba, K., Shia, J., Wang, J.Y., and Roehrl, M.H.A. (2022) Tumor stromal nicotinamide N-methyltransferase overexpression as a prognostic biomarker for poor clinical outcome in early-stage colorectal cancer. Sci Rep. 12, 2767

(18) Zhang, W., Rong, G., Gu, J., Fan, C., Guo, T., Jiang, T., Deng, W., Xie, J., Su, Z., Yu, Q., Mai, J., Zheng, R., Chen, X., Tang, X., and Zhang, J. (2022) Nicotinamide N-methyltransferase ameliorates renal fibrosis by its metabolite 1-methylnicotinamide inhibiting the TGF-beta1/Smad3 pathway. FASEB J. 36, e22084

(19) Wu, J.C., Merlino, G., and Fausto, N. (1994) Establishment and characterization of differentiated, nontransformed hepatocyte cell lines derived from mice transgenic for transforming growth factor alpha. Proc Natl Acad Sci U S A. 91, 674–678

(20) Varela-Rey, M., Martinez-Lopez, N., Fernandez-Ramos, D., Embade, N., Calvisi, D.F., Woodhoo, A., Rodriguez, J., Fraga, M.F., Julve, J., Rodriguez-Millan, E., Frades, I., Torres, L., Luka, Z., Wagner, C., Esteller, M., Lu, S.C., Martinez-Chantar, M.L., and Mato, J.M. (2010) Fatty liver and fibrosis in glycine N-methyltransferase knockout mice is prevented by nicotinamide. Hepatology. 52, 105–114

(21) Yen, C.L., Stone, S.J., Koliwad, S., Harris, C., and Farese, R.V., Jr. (2008) Thematic review series: glycerolipids. DGAT enzymes and triacylglycerol biosynthesis. J Lipid Res. 49, 2283–2301

(22) Eberle, D., Hegarty, B., Bossard, P., Ferre, P., and Foufelle, F. (2004) SREBP transcription factors: master regulators of lipid homeostasis. Biochimie. 86, 839–848

(23) Joshi, J.G., and Handler, P. (1960) Biosynthesis of trigonelline. J Biol Chem. 235, 2981–2983

(24) Song, Q., Chen, Y., Wang, J., Hao, L., Huang, C., Griffiths, A., Sun, Z., Zhou, Z., and Song, Z. (2020) ER stress-induced upregulation of NNMT contributes to alcohol-related fatty liver development. J Hepatol. 73, 783–793

(25) Bligh, E.G., and Dyer, W.J. (1959) A rapid method of total lipid extraction and purification. Can J Biochem Physiol. 37, 911–917

(26) Izumi, Y., Matsuda, F., Hirayama, A., Ikeda, K., Kita, Y., Horie, K., Saigusa, D., Saito, K., Sawada, Y., Nakanishi, H., Okahashi, N., Takahashi, M., Nakao, M., Hata, K., Hoshi, Y., Morihara, M., Tanabe, K., Bamba, T., and Oda, Y. (2019) Inter-Laboratory Comparison of Metabolite Measurements for Metabolomics Data Integration. Metabolites. 9,

(27) Fushimi, T., Izumi, Y., Takahashi, M., Hata, K., Murano, Y., and Bamba, T. (2020) Dynamic Metabolome Analysis Reveals the Metabolic Fate of Medium-Chain Fatty Acids in AML12 Cells. J Agric Food Chem. 68, 11997–12010

(28) Kim, D., Paggi, J.M., Park, C., Bennett, C., and Salzberg, S.L. (2019) Graph-based genome alignment and genotyping with HISAT2 and HISAT-genotype. Nat Biotechnol. 37, 907–915

(29) Danecek, P., Bonfield, J.K., Liddle, J., Marshall, J., Ohan, V., Pollard, M.O., Whitwham, A., Keane, T., McCarthy, S.A., Davies, R.M., and Li, H. (2021) Twelve years of SAMtools and BCFtools. Gigascience. 10,

(30) Liao, Y., Smyth, G.K., and Shi, W. (2014) featureCounts: an efficient general purpose program for assigning sequence reads to genomic features. Bioinformatics. 30, 923–930

(31) Chen, S., Zhou, Y., Chen, Y., and Gu, J. (2018) fastp: an ultra-fast all-in-one FASTQ preprocessor. Bioinformatics. 34, i884–i890

(32) Reimand, J., Arak, T., Adler, P., Kolberg, L., Reisberg, S., Peterson, H., and Vilo, J. (2016) g:Profiler-a web server for functional interpretation of gene lists (2016 update). Nucleic Acids Res. 44, W83–89

